# Human stem cell model of neural crest cell differentiation reveals a requirement of SF3B4 in survival, maintenance, and differentiation

**DOI:** 10.1101/2024.01.25.577202

**Authors:** Casey Griffin, Jean-Pierre Saint-Jeannet

**Author notes:** Corresponding author: Jean-Pierre Saint-Jeannet.

## Abstract

*In vitro* modeling is a powerful approach to investigate the pathomechanisms driving human congenital conditions. Here we use human embryonic stem cells (hESCs) to model Nager and Rodriguez syndromes, two craniofacial conditions characterized by hypoplastic neural crest-derived craniofacial bones, caused by pathogenic variants of SF3B4, a core component of the spliceosome. We observed that siRNA-mediated knockdown of *SF3B4* interferes with the production of hESC-derived neural crest cells, as seen by a marked reduction in neural crest gene expression. This phenotype is associated with an increase in neural crest cell apoptosis and premature neuronal differentiation. Altogether these results point at a role of SF3B4 in neural crest cell survival, maintenance, and differentiation. We propose that the dysregulation of these processes may contribute to Nager/Rodriguez syndrome associated craniofacial defects.

## Introduction

The *SF3B4* gene encodes the protein SAP49, which is a component of the U2 subunit of the major spliceosome (Will and Luhrmann, 2011). During pre-mRNA splicing, SAP49 binds upstream of the branch point sequence, tethering the U2 complex to the branch site, and allowing for proper splice site recognition and recruitment of the active subunits (Champion-Arnaud and Reed, 1994).

SF3B4 plays numerous roles within the cell, indicating importance beyond the spliceosome (Xiong and Li, 2020). Studies in *Xenopus laevis* and in humans provided evidence of transcriptional regulation by SF3B4 (Devotta et al., 2016; Marques et al., 2016), while studies in *Arabidopsis* have shown that SF3B4 can recruit RNA polymerase II complex to certain genes during embryogenesis (Xiong et al., 2019). SF3B4 has also been involved in controlling the translation as a co-factor at the endoplasmic reticulum, regulating biosynthesis of secreted proteins (Ueno et al., 2019). Finally, SF3B4 has been implicated as having a role in cell signaling by interacting with and controlling the levels of BMPR-IA at the cell surface (Nishanian and Waldman, 2004; Watanabe et al., 2007).

SAP49/SF3B4 variants in humans are associated with Nager and Rodriguez syndromes (OMIM#154400 and OMIM#201170), two rare forms of acrofacial dysostosis in which the craniofacial and appendicular skeleton are hypoplastic, with varying severity. Interestingly, in these patients the craniofacial skeletal structures affected are primarily derived from the neural crest (NC). The NC is a developmentally transient population of highly migratory cells that develop from the neural plate border, delaminate, and undergo an epithelial-to-mesenchymal transition, and populate the pharyngeal arches of the developing embryo. NC give rise to multiple cell types, including neurons, glia, smooth muscle cells melanocytes, muscle, and in the head form most of the craniofacial skeleton. While it has been shown that haploinsufficiency of SF3B4 causes Nager/Rodriguez syndrome, the mechanism by which a mutation in this component of the spliceosome leads to a cell-type specific disorder is unknown.

Here we use human embryonic stem cells (hESCs) to model the role of SF3B4 during NC cell (NCC) differentiation to gain insights into the pathomechanisms driving Nager/Rodriguez syndrome. We show that siRNA-mediated knockdown of SF3B4 leads to decreased NCC formation, increased apoptosis, and precocious neuronal formation. These results indicate that SF3B4 is necessary for NCC survival, maintenance, and differentiation, the disruption of these activities may therefore contribute to the craniofacial defects observed in Nager/Rodiguez syndrome patients.

## Results

### Human *in vitro* model of NCC differentiation

To study the role of SF3B4 during NCC differentiation, we turned to a well-established protocol of *in vitro* NCC derivation from hESCs (Fig. 1A) (Bajpai et al., 2010). Briefly, hESCs are plated in a defined differentiation medium and within 24 hours, the hESCs begin to form neuroectodermal spheres. These spheres will eventually settle at the bottom of the dish around day 5-7. Within 24-48 hours of settling, the spheres give rise to migrating NCC. By day 12 a majority of the spheres have settled and given rise to NCC, and washing away the spheres allows for a pure population of NCC. Prior to differentiation, hESCs express the pluripotency factor SOX2, and the epithelial marker E-CADHERIN (Fig. 1B). As differentiation proceeds, SOX2 and E-CADHERIN expression progressively decreases, while the expression of NCC genes is activated, until there is a pure population of NCC expressing both TFAP2A and P75/NGFR (Fig. 1C) as previously reported (Bajpai et al., 2010). This differentiation protocol is technically simple because differentiation can be monitored phenotypically by light microscopy, with the spheres and migrating NCC clearly distinguishable in the dish throughout the process (Fig. 1D).

**Figure 1:**
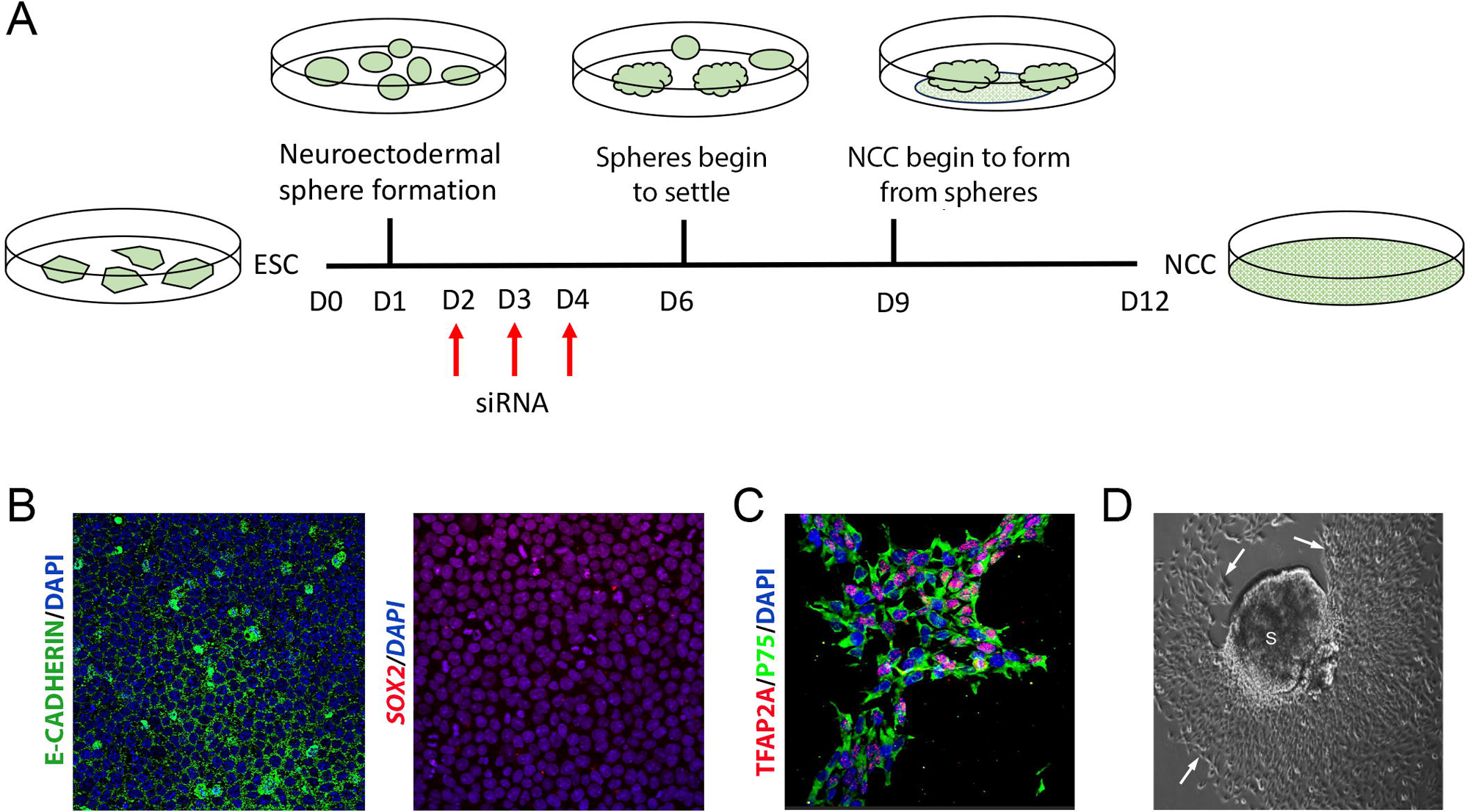
Human *in vitro* model of NCC differentiation. (**A**) Schematic diagram of the NCC differentiation protocol. Red arrows indicate timepoints of siRNA addition to the culture medium. (**B**) Confocal images of hESCs stained for E-CADHERIN (green) and SOX2 (red). (**C**) Confocal image of hESC-derived NCC stained for TFAP2A (red) and P75/NGFR (green) at day 12. (B,C) DAPI (blue) is used for nuclear staining. (**D**) Brightfield image of neuroectodermal sphere (S) with migrating NCC (arrows) at day 7.

### SF3B4 knockdown by siRNA in differentiating NCC

By qRT-PCR, SF3B4 is expressed during the entire process of NCC differentiation *in vitro*, with decreased expression levels toward the end of the differentiation protocol, around day 12 (Fig. 2A). To analyze the effect of knocking down SF3B4 on NCC differentiation, we used the siRNA technology. Pools of siRNA targeting SF3B4 or nontargeting controls were added to the cell culture medium on day 2-4 of the NCC differentiation protocol (Fig. 1A). Cells were collected for downstream analysis at day 4 (neuroectodermal spheres), day 7 (mixed population of spheres and NCC), or day 12 (NCC). Western blot analysis indicates that SF3B4 siRNA-mediated knockdown cause reduced SF3B4 expression as early as day 4 as compared to control siRNA treated samples, and by day 12 the expression levels of SF3B4 was completely eliminated (Fig. 2B). These results demonstrate that SF3B4 is expressed in hESC-derived NCC, and efficient silencing of SF3B4 can be achieved through siRNA-mediated knockdown.

**Figure 2:**
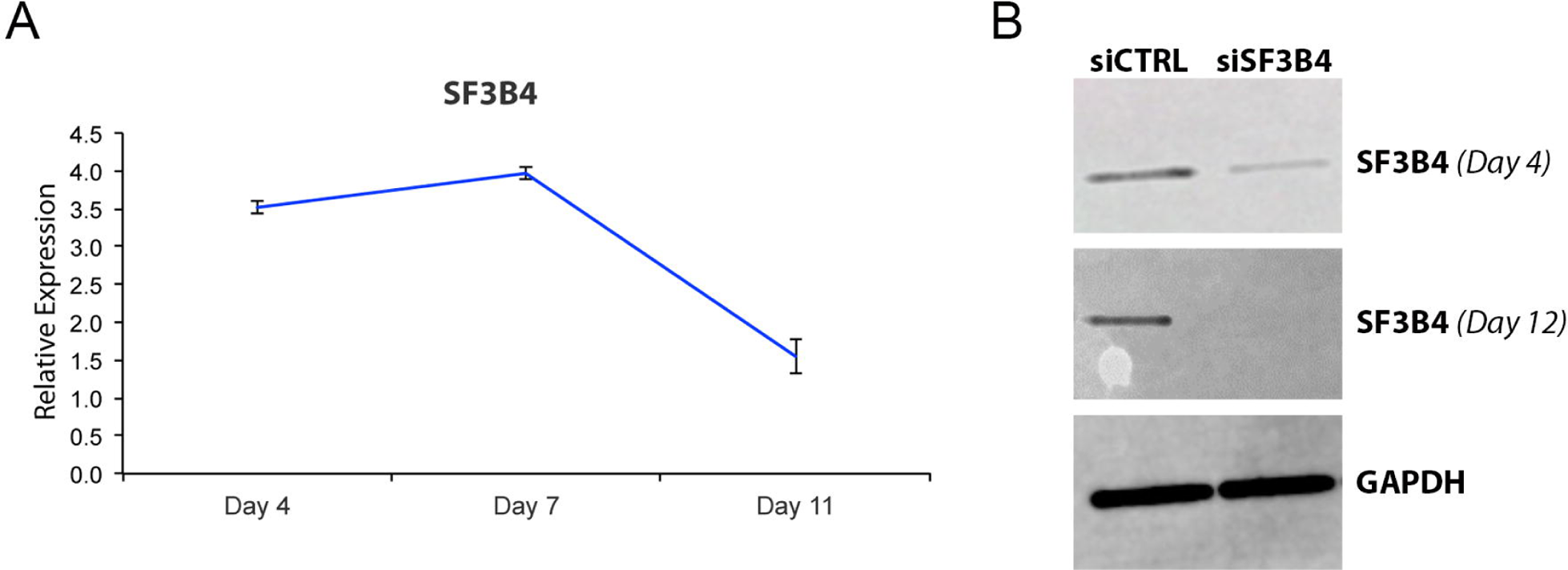
*SF3B4* expression and knockdown with siRNA. (**A**) qRP-PCR analysis of *SF3B4* expression during NCC differentiation *in vitro*. Values are normalized to *GAPDH* expression. (**B**) Western blot analysis showing SF3B4 expression levels at day 4 (D4) and day 12 (D12) of NCC differentiation after siRNA treatment. GAPDH is used as a loading control.

### Knockdown of SF3B4 affects neural crest formation

We next analyzed the impact of SF3B4 knockdown on the differentiation of hESC-derived NCC at day 7, and day 12 using Western blot, immunocytochemistry, and qRT-PCR. The neurotrophin receptor P75/NGFR and the HNK1 carbohydrate epitope are typically expressed by migrating NCC (Heuer et al., 1990; Bronner-Fraser, 1986), and both P75/NGFR and HNK1 expression levels were decreased upon SF3B4 knockdown as shown by Western blot at day 7 (Fig. 3A). This reduced expression was confirmed by immunofluorescence analysis at day 12 (Fig. 3B). Further, we used qRT-PCR to expand the range of NC markers analyzed to include two NC-specific genes, *TFAP2A* and *SOX10* (Rada-Iglesias et al., 2012; Pusch et al., 1998), in addition to *P75/NGFR* and *HNK1*. At day 7, expression levels of all 4 genes were significantly reduced in the SF3B4 siRNA samples as compared to control siRNA treated cells (Fig. 3C). These data suggest that SF3B4 is required for NC formation in this human *in vitro* model of NCC differentiation.

**Figure 3:**
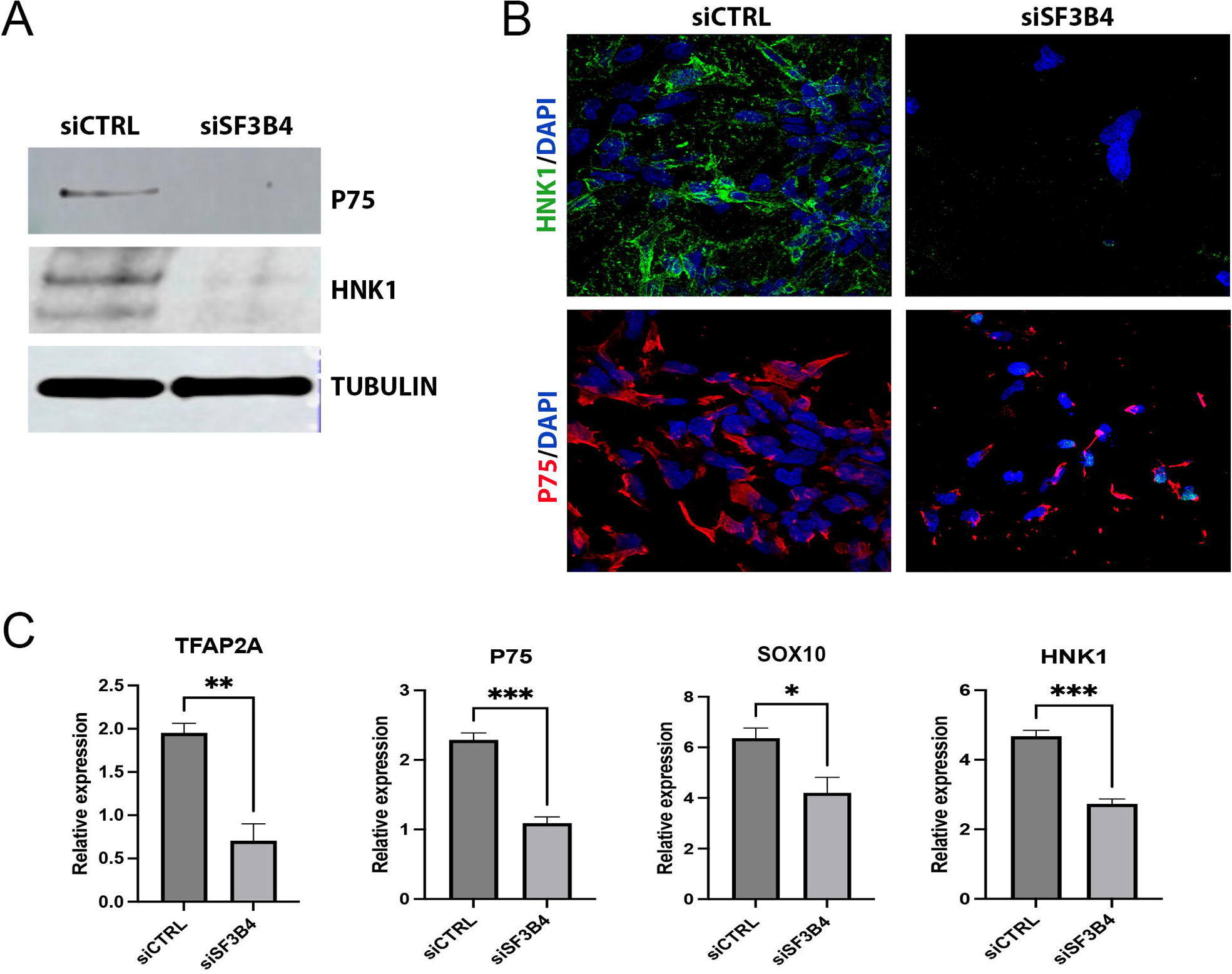
SF3B4 knockdown affects NCC formation. (**A**) Western blot analysis shows reduced P75/NGFR and HNK1 expression at day 7 of NCC differentiation upon siSF3B4 treatment as compared to controls (siCTRL). (**B**) Confocal images of NCC stained for HNK1 (green) and P75/NGFR (red) at day 12 of NCC differentiation with siRNA treatment (siSF3B4) compared to controls (siCTRL). DAPI (blue) is used for nuclear staining. (**C**) qRT-PCR analysis of *TFAP2A*, *P75/NGFR*, *SOX10*, and *HNK1* expression at day 12 of NCC differentiation, comparing siCTRL and siSF3B4 treatment. Values are normalized to *GAPDH* expression. * p<0.05, ** p<0.001, *** p<0.0001 (student’s unpaired t-test).

### Knockdown of SF3B4 causes precocious neuronal differentiation

During the NCC differentiation protocol, the number of NCC formed upon SF3B4 siRNA-mediated knockdown is greatly reduced as compared to control siRNA samples, as determined by the reduction in NCC genes expression (Fig. 3). We also noticed that the differentiated cells took on a neuronal morphology with extended neurite-like structures (Fig. 4A). In an attempt to characterize these cells, we performed qRT-PCR analysis for the expression of several neural genes including, PAX6, a marker of neural stem/progenitor cells (Sakurai and Osumi, 2008), GFAP, an intermediate filament expressed in astrocytes (Yang and Wang, 2015), NEUROD1, a transcription factor expressed by developing neurons (Lai et al., 2020), SOX2, a marker of multipotent neural stem cells (Ellis et al., 2004), DCX, a phosphoprotein typically expressed by immature neurons (Ma et al., 2010), HES1, a transcription factor that regulates maintenance and proliferation of neural stem cells (Ohtsuka and Kageyama, 2021), and VIMENTIN, an intermediate filament expressed by astrocytes and neural stem/progenitor cells (Yabe et al., 2003). While *PAX6*, *GFAP,* and *NEUROD1* were undetectable (not shown), we found that the expression of *HES1*, *VIMENTIN*, *DCX*, and *SOX2* was significantly increased upon SF3B4 knockdown as compared to control samples (Fig. 4B). These data suggest that SF3B4 is required for NCC maintenance, and with reduction of SF3B4 the remaining NCC adopt a neuronal fate.

**Figure 4:**
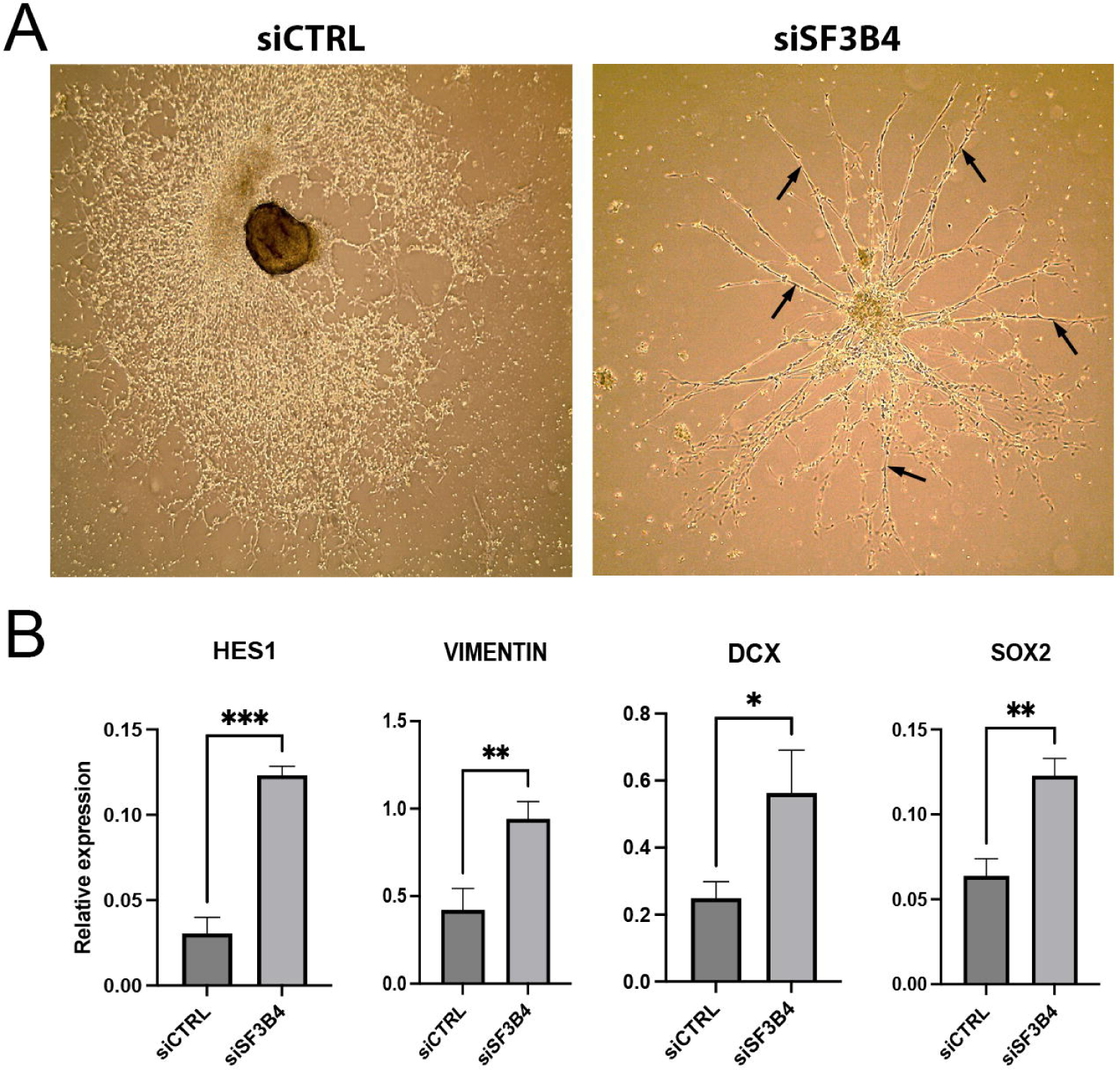
SF3B4 knockdown causes precocious neuronal differentiation. (**A**) Brightfield images of cells at day 12 of NCC differentiation showing the emergence of cells with extended neurite-like structures (black arrows) upon SF3B4 knockdown (siSF3B4) as compared to controls (siCTRL). (**B**) qRT-PCR analysis of *HES1, VIMENTIN, DCX* and SOX2 expression at day 7 of NCC differentiation. Values are normalized to *GAPDH* expression. * p<0.05, ** p<0.001, ns = no significance (student’s unpaired t-test).

### Knockdown of SF3B4 causes increased NCC apoptosis

We next wished to investigate the potential mechanism underlying the reduction in NCC observed upon SF3B4 knockdown during the differentiation protocol. At day 12, brightfield images showed large areas of dying cells in SF3B4 siRNA-treated samples as compared to control siRNA conditions, in which large fields of migrating NCC were observed (Fig. 5A). Furthermore, SF3B4 siRNA-treated neuroectodermal spheres at day 12 appeared unhealthy, with irregular edges and clumps of dying cells coming out of the spheres (Fig. 5B). We next analyzed the expression levels of P53, a factor associated with DNA repair and apoptosis, and several downstream effectors, PUMA and CASP3 (Audrey et al., 2018), to assess a possible increase in cell death in the SF3B4 siRNA treated samples. qRT-PCR analysis shows an early increase in *P53*, *PUMA*, and *CASP3* expression at day 4 of differentiation in SF3B4 depleted cells as compared to control cells (Fig. 5C). Western blot analysis also shows increased P53 expression at day 7 (Fig. 5D). Early activation of the P53 pathway is therefore the likely cause for the loss of cells observed upon SF3B4 knockdown, suggesting that SF3B4 is necessary for NCC survival.

**Figure 5:**
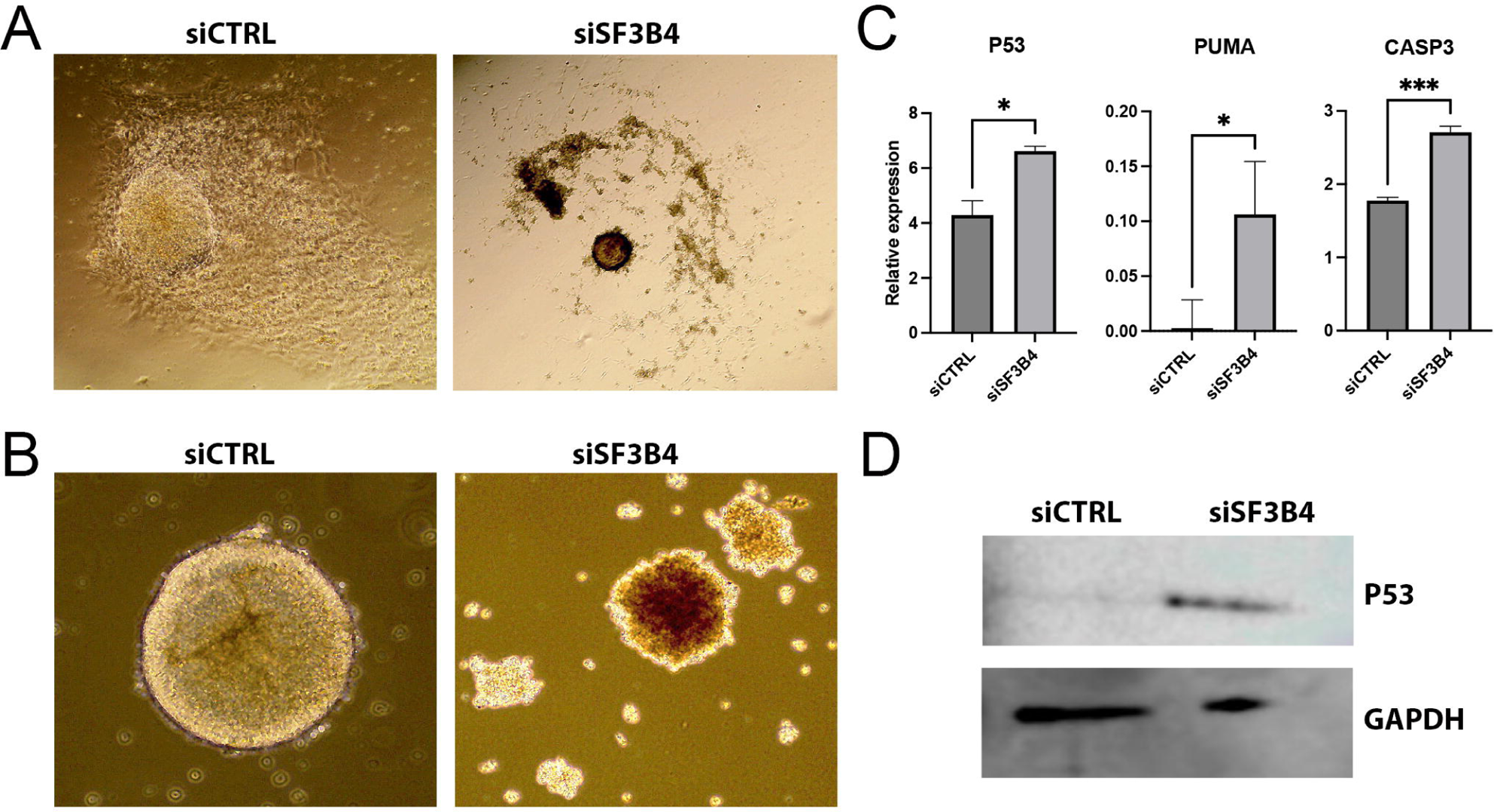
SF3B4 knockdown promotes NCC apoptosis. (**A**) Brightfield images of NCC differentiation at day 12 upon SF3B4 knockdown (siSF3B4) compared to control cells (siCTRL). (**B**) Brightfield images of spheres highlighting the spherical shape in controls (siCTRL) versus the irregular morphology with reduction of SF3B4 (siSF3B4). (**C**) qRT-PCR analysis of *P53*, *PUMA*, and *CASP3* expression at day 4 of NCC differentiation, comparing siCTRL and siSF3B4 treatments. Values are normalized to *GAPDH* expression. * p<0.05, ** p<0.001 (student’s unpaired t-test). (**D**) Western blot analysis showing increased P53 expression at day 7 of NCC differentiation in siSF3B4-treated cells versus control cells (siCTRL). GAPDH is used as a loading control.

## Discussion

Nager and Rodriguez syndromes are rare craniofacial spliceosomopathies characterized by hypoplastic NC-derived skeletal elements of the face, primarily caused by haploinsufficiency of SF3B4. Here we show that reduction in SF3B4 in a human *in vitro* model of NCC differentiation leads to a decrease in NCC formation through a mechanism that involves increased apoptosis, with remaining NCC adopting a neuronal fate. We propose that disruption of these processes is likely responsible for the presentation of the craniofacial phenotype in Nager/Rodriguez syndrome patients.

An interesting observation of this study suggests that a subset of cells that survive upon SF3B4 knockdown differentiate into neuron-like cells with long projections, which is not normally occurring under this NCC differentiation protocol – typically NCC self-renew and survive for a few weeks before dying off. *In vivo,* NCC give rise to various types of neurons and glia during development, with contributions to autonomic and sensory neuronal lineages. While the NCC generated under this *in vitro* differentiation protocol can also give rise to several types of neurons, their differentiation must be triggered by a change in differentiation medium. We show that neuronal differentiation is occurring with the loss of SF3B4, overriding the need for environmental cues to initiate neuron formation. It is unclear if the premature emergence of neuronal cells in condition of reduced SF3B4 activity is also occurring in human patients, and participate in the presentation of the craniofacial disease. One can speculate that this may indicate that these NCC differentiate into neurons at the expense of another NC-derived lineage essential to build the craniofacial skeleton, the chondrocytes. Future experiments will test whether SF3B4-depleted NCC are lineage biased, and have lost the ability to initiate a chondrocyte differentiation program.

Consistent throughout work aimed at understanding the mechanisms in various spliceosomopathies is the role of apoptosis. Other craniofacial spliceosomopathies have been linked to increased cell death in the head region. For example, EFTUD2, SNRPB, or TXNL4A knockdown in *Xenopus laevis* causes increased cell death resulting in abnormal craniofacial development (Park et al., 2022). Work in mouse has shown SNRPB and EFTUD2 mutations both leads to an increase in apoptosis (Alam et al., 2022; Beauchamp et al., 2021). Our work shows another clear connection between mutations in the spliceosome and increased apoptosis, emphasizing the importance of proper spliceosome function for NCC survival.

Previous studies in mouse, fish, and frog have strived to model Nager/Rodriguez syndrome in order to tease apart the underlying mechanism of the pathology. In mouse, it has been shown that homozygous *Sf3b4* mutants are embryonic lethal (Yamada et al., 2020), while heterozygous *Sf3b4* mutants have microcephaly and homeotic posteriorization of the vertebrae (Yamada et al., 2020; Kumar et al., 2023). The axial skeleton defects have been attributed to mis-splicing of chromatin remodelers and dysregulation of Hox gene expression (Kumar et al., 2023). In zebrafish, *sf3b4* mutation causes massive cell death in the retina of the eye (Ulhaq et al., 2023), reminiscent of another spliceosomopathy, retinitis pigmentosa (Griffin and Saint-Jeannet, 2020). In *Xenopus laevis*, morpholino-induced knockdown of Sf3b4 led to a reduction in NC gene expression resulting in hypoplastic craniofacial cartilages, attributed to an increase in NCC apoptosis (Devotta et al., 2016). In *Xenopus tropicalis,* CRISPR/Cas9-mediated Sf3b4 knockout showed that loss of one copy of the gene is dispensable for spliceosome activity. However, homozygous *sf3b4* mutant embryos exhibited reduced neural crest cell migration, increased apoptosis in the head region, reduced expression of *sox9*, a regulator of chondrogenesis, and ultimately death at the tadpole stage (Griffin, et al., 2024). While these studies have all been able to model different aspects of Nager/Rodriguez syndrome, here we have shown that the hESCs model of NCC differentiation can provide a unique perspective on the pathogenesis of this disorder and the cell-type specific activity of SF3B4.

Although siRNAs are powerful tools to evaluate gene function they also have limitations. One such limitation is the inability to accurately model patient mutations. Patients are haploinsufficient for SF3B4, but the siRNA represents a close to 100% knockdown of SF3B4 by day 12 of differentiation. An important next step in modeling Nager/Rodriguez syndrome is to develop patient-derived induced pluripotent stem cells (iPSCs) carrying the exact patient mutations in *SF3B4*. Patient-derived iPSCs allowing for a more precise characterization of the patient phenotype, and more rigorous transcriptomic and lineage bias analyses.

## Materials and Methods

### Human Embryonic Stem Cells and Their Maintenance

H9 (WA09, Wicell Research Institute, Madison, Wisconsin) human ESCs were used as wildtype hESCs with normal karyotype. hESCs were grown and expanded on growth-factor reduced geltrex-coated (Gibco, #A1413302) dishes in mTeSR^TM^1 medium (Stem Cell Technologies, #85850) to 80% confluence. Medium was changed daily, cells were split every 5-7 days with accutase (Stem Cell Technologies, #07920) (Bajpai et al., 2008) and plated at a 1:6 dilution on geltrex-coated plates.

### Neural Crest Cell Differentiation

For NCC differentiation, ESCs were collected using collagenase IV (Gibco, #17104-019), rinsed with PBS to remove all traces of enzyme, and transferred to serum-free NCC medium, as described (Bajpai et al., 2010). Medium was changed every 3 days.

### siRNA Knockdown

Accell siRNA targeting human SF3B4 (Cat No. E-017190-00-0050) and siControl (Cat No. D-001910-10-50) were purchased from Horizon Discovery (Boyertown, PA). Accell siRNAs do not require transfection and can be added directly to the culture medium. For siRNA knockdown, siRNA was added to the differentiation medium at days 2, 3, and 4 at a concentration of 330 nM. Differentiation was carried out until day 4, day 7, or day 12 and the cells collected for downstream analysis.

### Western Blot Analysis

Cells were collected in RIPA lysis buffer, and western blot analysis was performed as previously described (Devotta et al., 2016). Primary antibodies were as follows: anti Sf3b4 polyclonal antibody (Proteintech, Cat No. 10482-1-AP, 1:2000 dilution), anti NGFR/p75NTR polyclonal antibody (Sino Biological, Cat No. 102002-T34, 1:2000 dilution), anti p53 monoclonal antibody (Invitrogen, Cat No. MA1-12549, 1:1000 dilution), anti HNK-1/N-CAM monoclonal antibody (Sigma, Cat No. C6680-100TST, 1:1000 dilution). Secondary antibodies were anti-rabbit (EMD Millipore, Cat No. MAB201P) and anti-mouse (Abcam, Cat No. ab6820) IgG coupled to horseradish peroxidase (1:10,000 dilution)

### Immunocytochemistry

Cells were plated on prewashed glass coverslips coated with fibronectin and grown to 60-80% confluence (usually 2 days after plating). They were then fixed using fresh 4% formaldehyde. Cells were washed with PBS and place in PBTx blocking solution (1% BSA, 0.1% Triton-X in PBS) for 1 hour at room temperature. Primary antibodies were brought to the desired dilution (1:100 to 1:1000) in PBTx and incubated with cells overnight at 4°C (E-cadherin, Abcam, Cat No. ab231303; Sox2, Santa Cruz Biotechnologies, Cat No. sc-17320; TFAP2A, Abcam, Cat No. ab108311; P75/NGFR, Santa Cruz Biotechnologies, Cat No. sc-5634; HNK-1, Sigma, Cat No. C6680-100TST). Afterward, the cells were washed with PBTx. Fluorophore conjugated secondary antibodies diluted in PBTx (Invitrogen, Cat Nos. A32723, A32731, A32727, A32732, 1:1000) were incubated with cells for 45 minutes in the dark, at room temperature. Cells were washed again with PBS, then DAPI (0.1 mg/mL in PBS) was added for 30 seconds, before final washing with PBS. Coverslips were mounted on slides using 20% glycerol and imaged using a Leica SP8 confocal microscope. Images of five random fields were taken for each sample, images in figures are representative.

### Quantitative PCR Analysis

Confluent cells were collected using accutase and total RNA was extracted using RNeasy microRNA isolation kit (Qiagen, Cat No. 74004), with the samples digested with RNase-free DNase I to eliminate genomic DNA. RT-qPCR analysis was performed using Power SYBR Green RNA to CT 1 step RT-PCR kit (Applied Biosystems, #4389986) on a QuantStudio 3 Real-Time PCR System (Applied Biosystems, Foster City, CA). The primers were predesigned qPCR assays for human genes (IDT, Coralville, IA) as follows: GAPDH (Assay ID: Hs.PT.39a.22214836), TFAP2A (Assay ID: Hs.PT.58.5602), SOX10 (Assay ID: Hs.PT.58.4891394), P75/NGFR (Assay ID: Hs.PT.58.4045496), HNK1/B3GAT1 (Assay ID: Hs.PT.58.2273334), TP53 (Assay ID: Hs.PT.58.39676686), PUMA (Assay ID: Hs.PT.58.39966045), CASP3 (Assay ID: Hs.PT.56a.25882379.g), SOX2 (Assay ID: Hs.PT.58.237897.g), DCX (Assay ID: Hs.PT.58.118505), PAX6 (Assay ID: Hs.PT.58.25914558), GFAP (Assay ID: Hs.PT.58.14980282), NEUROD1 (Assay ID: Hs.PT.58.38524795), HES1 (Assay ID: Hs.PT.58.4181121), VIMENTIN/VIM (Hs.PT.58.38906895).

## Acknowledgements

We thank members of the Saint-Jeannet laboratory past and present for their support and helpful discussions.

## Funding

This work was supported by grants from the National Institutes of Health to J-P.S-J. (R01-DE025468) and C.G. (F32-DE030699).

## Competing Interests

The authors declare no competing interests.

## Notes

### Competing Interest Statement

The authors have declared no competing interest.

### Summary of Updates

Updated Figure 4 and Figure 5 to clarify results.

